# Interactions between SARS-CoV-2 N-protein and α-synuclein accelerate amyloid formation

**DOI:** 10.1101/2021.04.12.439549

**Authors:** Slav A. Semerdzhiev, Mohammad A. A. Fakhree, Ine Segers-Nolten, Christian Blum, Mireille M. A. E. Claessens

## Abstract

First cases that point at a correlation between SARS-CoV-2 infections and the development of Parkinson’s disease have been reported. Currently it is unclear if there also is a direct causal link between these diseases. To obtain first insights into a possible molecular relation between viral infections and the aggregation of α-synuclein protein into amyloid fibrils characteristic for Parkinson’s disease, we investigated the effect of the presence of SARS-CoV-2 proteins on α-synuclein aggregation. We show, in test tube experiments, that SARS-CoV-2 S-protein has no effect on α-synuclein aggregation while SARS-CoV-2 N-protein considerably speeds up the aggregation process. We observe the formation of multi-protein complexes, and eventually amyloid fibrils. Microinjection of N-protein in SHSY-5Y cells disturbed the α-synuclein proteostasis and increased cell death. Our results point toward direct interactions between the N-protein of SARS-CoV-2 and α-synuclein as molecular basis for the observed coincidence between SARS-CoV-2 infections and Parkinsonism.

## Introduction

Symptoms of SARS-CoV-2 infections that cause the current Covid-19 pandemic are not limited to the respiratory tract. The virus also affects other organs and tissues. SARS-CoV-2 has been found in neurons in different brain regions.^1, 2^ For many of the patients infected with SARS-CoV-2, acute and subacute neurological complications have been reported.^3-5^ One of these complications, the loss of smell, is a common premotor symptom in Parkinson’s disease (PD). This symptom and the recent reports of cases of Parkinson’s disease in relatively young patients after a SARS-CoV-2 infection suggests that there may be a link between SARS-CoV2-infections and the development of PD.^6^

The link between viral infections and neurodegeneration is established for some viruses.^7-10^ The most well-known example is the 1918 influenza pandemic (Spanish flu) which coincided with an increase of encephalitis lethargica, followed by numerous cases of post-encephalitic Parkinsonism.^11, 12^ In more recent times, multiple indications of a relation between PD and viral infections have been reported.^10, 13^ Whether viral infections indirectly cause neurodegeneration via the immune system^14^ or if the effect is direct is unclear. Neurodegenerative diseases such as Alzheimer’s Disease (AD) and PD are protein aggregation diseases in which specific proteins, tau and Aβ peptide in AD and α-synuclein (αS) in PD, assemble into amyloid aggregates. Once started, the aggregation process spreads from cell to cell and the formed aggregates and deposits hamper brain function.^15-17^ In a direct mechanism the virus itself triggers the protein aggregation process. The virus would thus be responsible for the onset of the pathological protein aggregation process. Indeed, such a direct relation has been found for Aβ peptide aggregation (AD) in model cell lines and animals infected with herpes simplex and respiratory syncytial virus.^18^

In light of the potential relation between SARS-CoV-2 infections and the development of Parkinson’s disease we investigate the direct effect of SARS-CoV-2 proteins on αS aggregation and αS proteostasis in model systems. We show, in test tube experiments, that SARS-Cov-2 S-protein has no effect on αS aggregation while SARS-CoV-2 N-protein considerably speeds up the aggregation process. N-protein and αS directly interact and this interaction results in the formation of complexes that contain multiple proteins, and eventually amyloid fibrils. Microinjection of N-protein in SHSY-5Y cells disturbed the αS proteostasis and increased cell death. Our results suggest that the observed link between SARS-CoV-2 infection and Parkinson’s disease might originate from a molecular interaction between virus proteins and αS.

## Results

The nucleocapsid protein (N-protein) and the spike protein (S-protein) are the most abundant, (partly) soluble structural SARS-CoV-2 proteins with copy numbers of ∼1000 and ∼300 monomers per virus particle respectively.^19^ The net positively charged N-protein packs the negatively charged viral genome into a higher order structures.^20^ The N-protein been assigned additional functions during viral infection.^21^ The S-protein is anchored to the membrane where it is exposed on the virus surface and plays a role in receptor recognition, docking and virus entry.^22, 23^ The S-protein is the main target in vaccination strategies since it induces the immune response of the infected host. The N-protein is also considered as a target for vaccine development because in the SARS family of viruses the N-protein gene is more conserved and stable than the S-protein gene.^24, 25^

Considering their abundance in infected cells we assess if interactions between αS and N-protein or S-protein affect the aggregation of αS into amyloid fibrils in *in vitro* experiments. In these experiments we followed the aggregation of αS in the presence and absence of N-protein and S-protein using Thioflavin T (ThT) fluorescence assays. The fluorescence of the dye ThT increases upon binding to amyloid fibrils. ThT fluorescence can therefore be used as a direct readout for fibril formation. A key parameter in such assays is the time to the first (visible) onset of aggregation or aggregation lag time. This time reflects how fast aggregation prone nuclei appear and amyloid fibrils are formed. It thus quantifies the impact of external triggers, like additional proteins, on the aggregation of αS. In the absence of additional proteins, the onset of aggregation of αS is observed at time scales > 240 hours (Fig. 1a). We observe no change in the outcome of the aggregation assay in the presence of S-protein (Fig. 1a). In the presence of N-protein we see a strong decrease in the time to the onset of aggregation which reduces to < 24 hours (Fig. 1b). On long time scales a transition to a second plateau is observed in the presence of N-protein as discussed later in the text.

**Figure 1.**
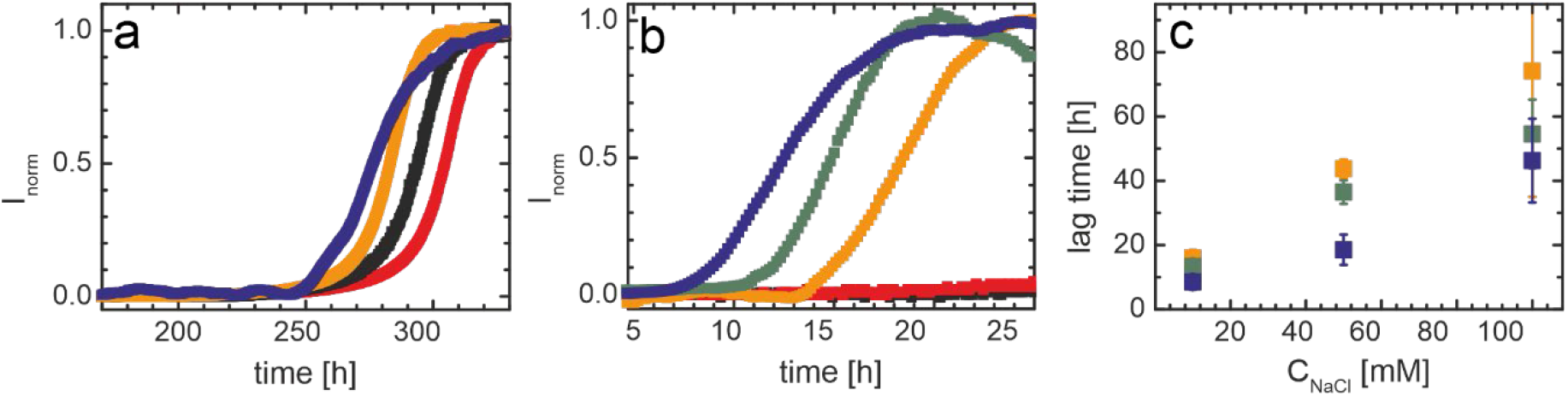
Aggregation of αS in the absence and presence of SARS-CoV-2 proteins. **a)** Aggregation assay of αS in the absence (black) and presence (color) of S-protein. The aggregation process is followed by recording the fluorescence of the amyloid binding dye ThT. The assay was performed at a NaCl concentration of 10 mM with 50 µM αS and 0.1 µM (red), 0.5 µM (orange) and 1 µM (blue) S-protein. The ThT fluorescence intensity (I) is normalized to the plateau value. **b**) ThT based aggregation assay of αS in the presence of N-protein. The assay was performed at a salt concentration of 10 mM NaCl with 50 µM αS and 0 µM (black) 0.1 µM (red), 0.5 µM (orange), 0.8 µM (green) and 1 µM (blue) N-protein. The ThT intensity (I) is normalized to the initial plateau value. **c)** Influence of the salt concentration on aggregation lag time for N-protein concentrations of 0.5 µM (orange), 0.8 µM (green) and 1 µM (blue) at an αS concentration of 50 µM. The points represent the mean of 3 independent measurements and error bars show the standard deviation.

We exclude that the increase in ThT fluorescence is the result of N-protein aggregation; at identical concentrations, incubation of N-protein alone does not increase the ThT fluorescence intensity (See SI Figure S1). To quantify the change in the onset of αS aggregation we determine the aggregation lag time. With increasing N-protein concentration the αS aggregation lag time decreases. This concentration dependent decrease evidences that direct interactions between αS and N-protein trigger αS aggregation.

N-protein and αS are net oppositely charged, near neutral pH (7.4) the calculated net charge amounts to +24e and –9e respectively. Electrostatic attraction is therefore likely to play a role in the inter-molecular interactions. Increasing the ionic strength of the solution, and thus screening the electrostatic charge, indeed increases the aggregation lag time (Fig 1c). However, also at higher salt conditions the lag time is still considerably shorter in the presence N-protein compared to the control. Moreover, the N-protein concentration dependence of the decrease in the lag time is conserved. Even at the highest salt concentration tested, the time scales at which we observe αS aggregation do not revert to the > 240 hours observed for αS alone. We therefore conclude that besides electrostatics other attractive forces contribute to the interaction between αS and N-protein.

To obtain insight into the strength of the interaction between αS and N-protein we performed a microscale thermophoresis assay. Microscale thermophoresis (MST) relies on changes in the diffusion coefficient of a particle upon binding to a partner. In the experiment one of the partners needs to be fluorescently labelled and labelling the one with the lower molecular weight provides optimal contrast. In our experiments we therefore fluorescently labelled αS. The concentration of fluorescently labelled αS was kept constant and the MST response was studied as a function of the N-protein concentration. The lower and higher plateau in the MST signal were interpreted as free αS and αS bound to N-protein respectively (Figure 2a). We observe a sharp transition from the unbound to bound state of αS and an EC_50_ of approximately 0.3 µM. The steepness of the transition indicates that binding is cooperative.

**Figure 2.**
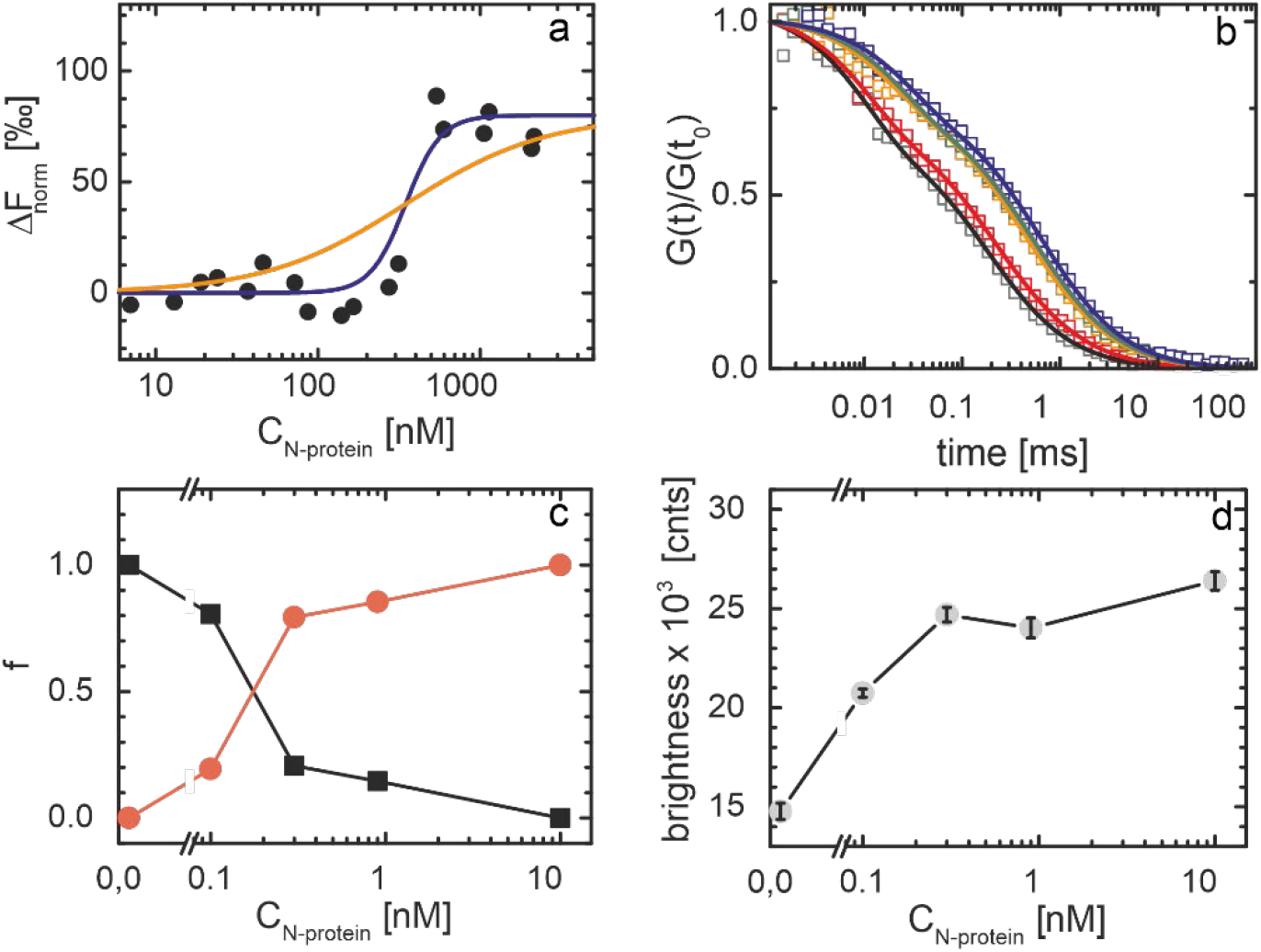
Interaction between αS and N-protein. a) Binding curve characterizing the interaction between αS and N-protein obtained from microscale thermophoresis (MST) experiments. MST data points are shown in black. The lines indicate binding with a Hill coefficient of 1 (orange) and 4 (blue) and an EC_50_ of 0.3 µM. These curves serve as a guide to the eye. b) FCS autocorrelation curves normalized to G(t_0_) (where t_0_ = 0.001 ms) (hollow symbols) of fluorescently labelled αS in the presence of 0 µM (black), 0.1 µM (red), 0.3 µM (orange), 0.9 µM (green) and 10 µM (blue) N-protein. Not all data points are shown for clarity. Fits to the autocorrelation curves are visible as lines with the corresponding color. In fitting the curves a triplet fraction was considered, and a fast and a slow diffusing component when necessary. c) Fractions (f) of the slow (red) and fast (black) diffusing components obtained from the fits to the autocorrelation curves shown in b. The errors bars show the standard deviation in the observed brightness. d) Average particle brightness as a function of the N-protein concentration.

Fluorescence correlation spectroscopy (FCS) experiments were performed to obtain first insights into the number of αS molecules in αS/N-protein complexes. In the FCS experiment αS was fluorescently labelled. Upon increasing the N-protein concentration we observe a strong shift in the correlation curves to longer times, indicating the formation of slow diffusing complexes (Figure 2b). For αS alone, the FCS autocorrelation curve can be fitted to a single diffusing species with a diffusion coefficient of approximately 86 µm^2^/s, in agreement with the expected size of the protein and earlier findings.^26^ With increasing N-protein concentration a second slower diffusing species with a diffusion coefficient of approximately 27 µm^2^/s appears. In figure 2c we plot the fraction of both the slow and fast diffusing species as a function of the N-protein concentration. In agreement with the MST data, we observe a transition from the unbound to bound state that is cooperative and an EC_50_ of ∼0.2 µM. Concomitant with the appearance of the αS/N-protein complex, we observe an increase of the average brightness of the diffusing protein complexes (Figure 2d). In the αS/N-protein complex the brightness increased by a factor of 1.8 times compared to unbound αS. Considering that of the total αS concentration of 20 nM, only half was labelled and ignoring the possible fluorescence quenching due to fluorophore-fluorophore interactions that has been observed in other protein systems under certain conditions,this indicates that on average 3 to 4 αS proteins are present in an αS/N-protein complex. Note that although this indicates that N-proteins accumulate αS, the FCS experiment was performed in excess of N-protein. ^27^ The aggregation experiments were performed in excess of αS, accumulation of even higher numbers of αS on N-protein is therefore likely. We conclude that the decrease in αS aggregation lag-time in the presence of N-protein results from direct interactions between the two protein species and the accumulation of αS on N-proteins.

The increase in ThT fluorescence intensity during αS aggregation in the presence of N-protein deviates from the typically observed pattern. Instead of the typical single step nucleation and growth process, we observe two growth steps. At longer time scales a second, higher, plateau in the ThT fluorescence intensity appears (Figure 3a). To verify that in both plateaus the observed ThT fluorescence results from the formation of amyloid fibrils, we performed atomic force microscopy (AFM) experiments. We obtained samples for AFM experiments that cover both the first and the second plateau in ThT fluorescence. AFM images obtained from samples in both plateaus in ThT fluorescence show the presence of helical amyloid fibrils (Figure 3b,c). The AFM images however indicate that the morphology of the fibrils is different in both plateaus. In the first plateau, fibrils of two different morphologies can be discriminated. These fibrils differ in helical periodicity. To determine the periodicity of the helix, a direct Fourier transform (DFT) was performed on the images of fibrils formed in both plateaus. We observe two clear peaks in the DFT analysis at periodicities of approximately ≈130 nm and ≈90 nm (Figure 3d). The morphology of fibrils in the samples obtained in the second plateau is homogeneous (Figure 3d). DFT analysis shows that these fibrils have a periodicity of 130 nm (Figure 3d). This periodicity agrees with previous AFM and cryo-electron microscopy studies on the structure of αS amyloid fibrils in the absence of N-protein.^28^ N-protein speeds up the formation of αS fibrils but does not change the morphology of fibrils observed after long time incubation.

**Figure 3.**
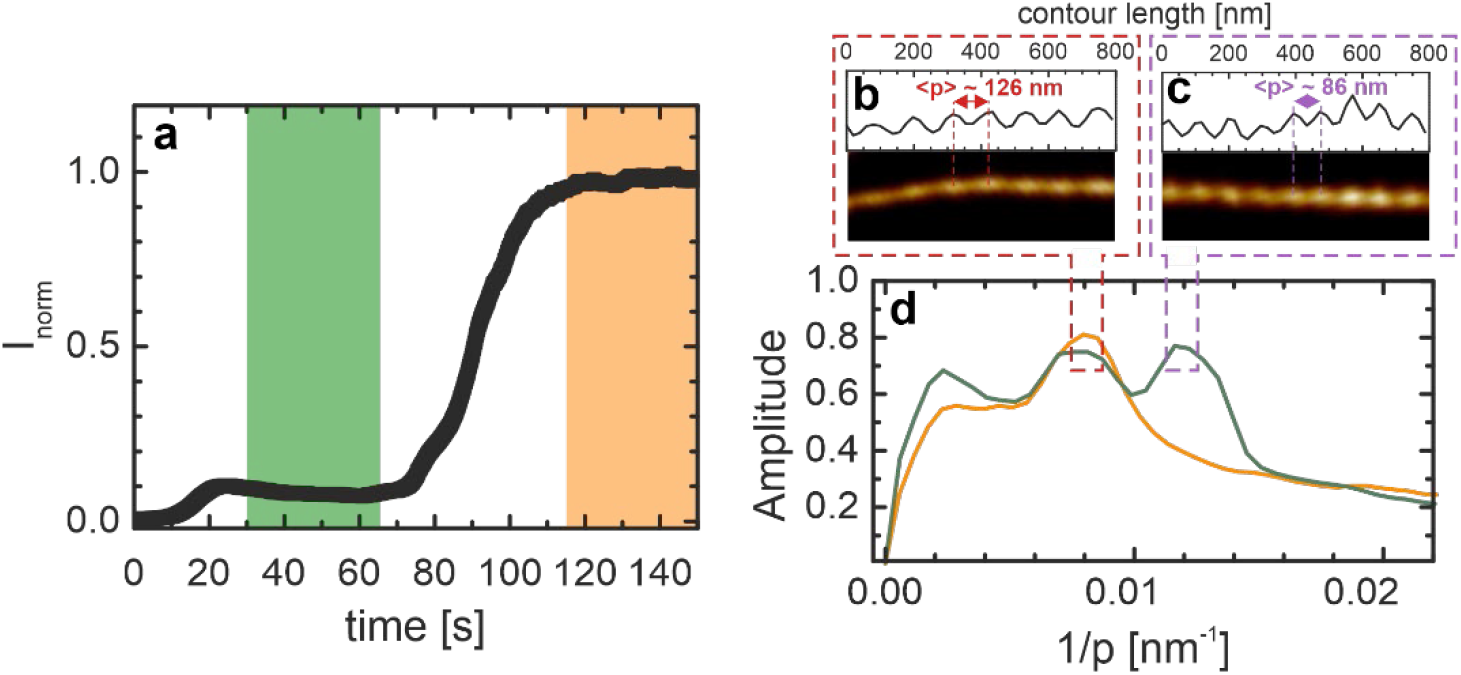
Aggregation of αS into amyloid fibrils in the presence of N-protein. a) Full ThT aggregation curve of αS in the presence of N-protein (1 µM N-protein, 50 µM αS, 50 mM NaCl). The data is normalized to the final plateau in ThT intensity. The colors refer to the color coding in d. b), c) In the AFM images two distinctly different fibril populations, with different morphologies, can be discriminated. The cross-sections along the length of the fibrils (top) show these fibrils differ in helical periodicity. d) Discrete Fourier analysis on fibrils obtained in the first (green) and second (orange) plateau. In first plateau 2 populations with different periodicity (*p)* of ≈90 nm and ≈130 nm are observed (green line). The peak at low 1/*p* values is an analysis artifact. In the second plateau, fibrils with a *p*=130 nm dominate the population, in the analysis fibrils with a periodicity *p*=90 nm are no longer visible (orange line). A total of 79 and 72 fibrils were analyze for the first and second plateau respectively.

Above we have shown proof that the direct interaction between N-protein and αS triggers αS aggregation into amyloid fibrils in *in vitro* experiments. Next, we have conducted microinjection experiments to investigate the effect of the presence of N-protein in a cellular context. From literature data we estimate the concentration of N-protein in infected cells to be of the order of 500 nM.^19^ The injected N-protein concentration and injection volume were chosen to approximately result in this concentration in the cells and to mimic concentrations expected in infections (see Materials and Methods).

The intrinsically disordered protein αS has been suggested to have many functions. In the cell, αS is found on trafficking vesicles and its main function most likely involves membrane remodeling in membrane trafficking processes.^29-36^ Bound to the membranes of vesicles αS adopts an α-helical conformation that can be discriminated from the unstructured protein (or other conformational states) in Förster resonance energy transfer (FRET) experiments.^37, 38^ These FRET probes used in *in vitro* experiments have also been applied to identify and localize membrane bound αS in cells.^39^ It is thus possible to use FRET to discriminate between conformational subensembles that potentially represent different functions and thereby gain insights into the αS proteostasis.

SH-SY5Y cells express αS and are a well-established neuronal cell model in PD research. With these cells, two different sets of experiments were performed. In one set, single cells were microinjected with both FRET labelled αS and N-protein and fixed 5 days after injection. In a second set of experiments the cells were injected with FRET labelled αS and N-protein and additional unlabelled αS and fixed 3 days after injection. We expect that the redistribution of αS from functional to dysfunctional states is a slow process. We hypothesize that by either giving the cells more time or by increasing the αS concentration this redistribution may become visible. In these experiments, the cells injected with only FRET labeled αS served as a control. After fixation, cells were counterstained with DAPI and imaged using a confocal fluorescence microscope.

Compared to the control where approximately 10% rounded up/dead cells are observed, we find approximately double the amount of dead cells after microinjection of N-protein. Typical images of the surviving injected cells are presented in Figure 4a. In the control, we see the previously reported distribution of αS between a high FRET vesicle bound and a low FRET cytosolic αS population.^39^ Vesicle bound αS is clearly present in orange (high FRET) fluorescent puncta, while cytosolic αS is visible as a spread out green (low FRET) background. The overall appearance of the FRET signal from cells that were co-injected with N-protein is very similar, both high FRET fluorescent puncta and low FRET spread out cytosolic signal can be found. However, in the cells that were microinjected with N-protein, we observe less high FRET (orange) signal compared to the control group (Fig. 4a). This indicates that in the presence of N-protein, the αS proteostasis is disturbed resulting in less vesicle bound αS.

**Figure 4.**
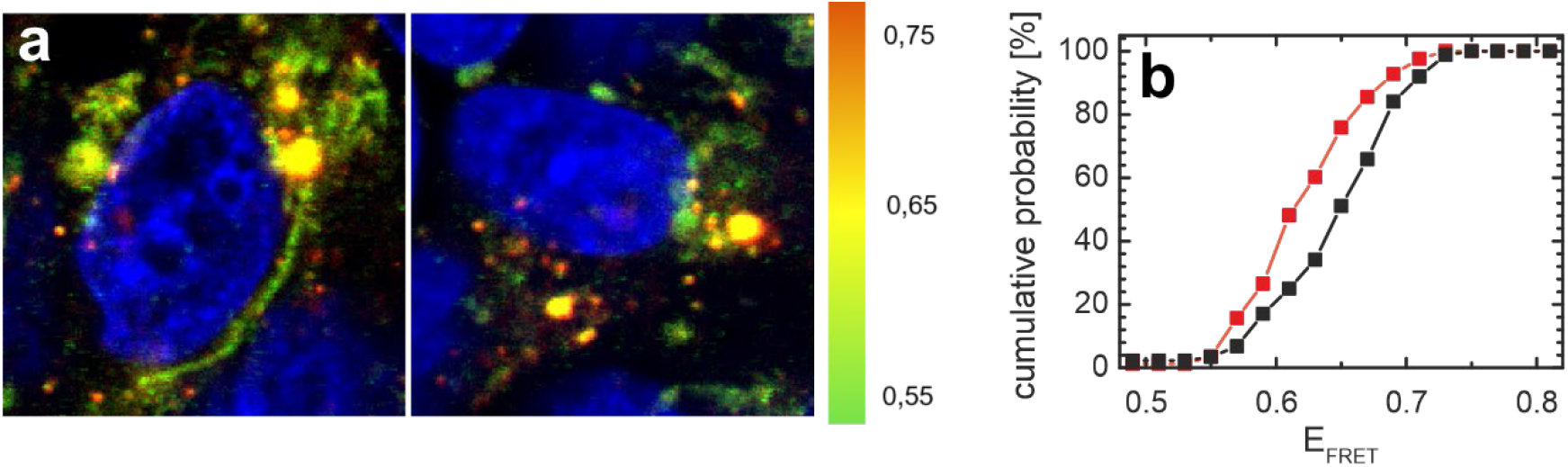
Distribution of αS in SH-SY5Y cells is affected by N-protein. **a**) FRET images of cells microinjected with the αS FRET probe. The color coding represents E_FRET_ (green: low E_FRET_; yellow: mid E_FRET_; orange: high E_FRET_). The cell nucleus is counterstained with DAPI and visible in blue. A representative image of cells co-injected with N-protein and the αS FRET probe is shown on the left, the control cells on the right were only injected with the αS FRET-probe. **b)** Distribution of average FRET efficiencies of αS per image for all cells injected with N-protein (red) and control cells (black). The cumulative histograms contain data from at least 80 images for both the control and the N-protein injected samples. The average FRET efficiency of αS in cells injected with N-protein is shifted to lower E_FRET_ values.

To substantiate the visual impression that in the presence of N-protein an overall average lower FRET value is observed, we estimated the αS FRET efficiency (E_FRET_ = intensity acceptor/(intensity donor + intensity acceptor)) averaged over all pixels for each image made (Materials and Methods). Subsequently we plotted cumulative histograms for the averaged αS FRET efficiency per image for the N-protein injected cells together with the controls (see SI Figure S2). The histograms obtained for 3 and 5 days after microinjection agree well. For both sets of experiments the distribution of the αS FRET efficiencies for the N-protein injected cells is shifted to lower values compared to the control. For the control samples, the mean αS FRET efficiencies quantitatively agree (see SI Figure S2). The width of the distribution is slightly enhanced for the control data obtained 5 days after microinjection. The good agreement between the data sets justifies accumulation of the data. In figure 4b we show the cumulative distribution of FRET efficiencies of αS in all control cells and all cells injected with N-protein. Compared to the control, the FRET efficiency distribution of αS in cells injected with N-protein is systematically shifted to lower values. Our data shows that the presence of N-protein results in redistribution of αS between (dis)functional conformational states. *In vitro* experiments shows that the observed change in FRET efficiency does not result from a direct interaction between the N-terminus of αS and N-protein. Even in the presence of excess N-protein where we expect all αS to be bound, we do not observe a change in FRET efficiency of the FRET labelled αS compared to the control (see SI Figure S3). The N-protein does apparently not interact with αS in the FRET labelled N-terminal region or this binding does not induce measurable conformational changes in αS.

## Discussion

The *in vitro* experiments on recombinantly expressed proteins show that SARS-CoV-2 S-protein does not affect the aggregation of αS into amyloid fibrils. The SARS-CoV-2 N-protein however, very effectively decreases the time to the onset of αS aggregation. Additionally, the aggregation process in the presence of N-protein differs from the aggregation process of αS alone. In the presence of N-protein, aggregation of αS proceeds in two steps, represented by two plateaus in the ThT fluorescence. The analysis of AFM images of fibrils shows that two populations of fibrils with a helical periodicity of approximately 90 nm and 130 nm are present during the first plateau phase. Only a single population of fibrils with a helical periodicity of 130 nm is found in the second plateau. The fibrils with the smaller helical periodicity are no longer observed. Although aggregation of αS in the presence of N-protein proceeds in two steps, first fibril nucleation is fast. The first plateau in the ThT fluorescence is found at a relatively low ThT intensity which indicates that the produced fibril mass is not very high. The fibrils formed in the first phase trigger the formation of or conversion to a second, thermodynamically more stable, fibril polymorph. The higher stability of this polymorph results in an increase of fibril mass and a plateau of higher ThT intensity. Note that the timescales on which the second polymorph appears is still fast compared to amyloid fibril formation in the absence of N-protein.

The MST and FCS data show that there is a direct interaction between N-protein and αS. The binding of the αS to the net positively charged N-protein appears to be mediated by attractive electrostatic interactions. This indicates that interaction involves the negatively charged C-terminus of αS. The absence of a change in the FRET efficiency of the αS FRET probe confirms that the interaction with the N-protein is likely mediated via the C-terminal region or the aggregation prone central NAC region of αS. In solution, electrostatic repulsion between net negatively charged αS proteins prevents their aggregation. Charge compensation due to binding to N-protein, not only exposes the aggregation prone NAC region of αS but also eliminates electrostatic repulsion of other αS proteins. Our data also shows that the complexes contain multiple αS proteins. The close proximity of multiple αS proteins in the complex in an aggregation prone conformation potentially facilitates the formation of a nucleus that triggers further aggregation and thus decreases the time to the onset of aggregation.^40^

Even in the complex environment of the cell we see clear signs that the presence of the N-protein markedly affects αS proteostasis. In the cell, αS exists in at least two different conformational sub-ensembles. The presence of the N-protein results in a change of the FRET efficiency distribution and hence in either a change in the population of the sub-ensembles or in the appearance of a new conformation. In the microinjection experiments on cells, the concentration of N-protein is low compared to the concentration of αS. It is therefore rather remarkable that we can detect a clear shift in the distribution of the αS FRET efficiencies.

The presence of N-protein indirectly affects the FRET efficiency of the αS FRET probe by disturbing αS proteostasis. The affinity of αS for N-protein is of the same order of magnitude or higher as the affinity reported for other αS interactions.^41-43^ Therefore, the N-protein will compete with other αS binding partners for interactions inside the cell. For reliable cellular performance, protein interaction networks must be robust. Small changes and additional binding partners are therefore not expected to easily disturb interaction networks. We however do see a clear redistribution of αS over (dis)functional states even at the rather short time scales studied. The observed increase in the fraction of images containing dead cells, further supports the idea of an imbalance in the cells proteostasis after injection with N-protein. We cannot confirm that this imbalance is the result of αS aggregation and the presence of amyloid fibrils in the cells injected with N-protein although we observe elongated and fibril mesh like structures. Considering that Parkinson’s disease typically develops on very long time scales, absence of fibrils in the microinjected cells would not be surprising.

Summarizing, we have identified a SARS-CoV-2 protein that induces the aggregation of αS in the test tube. In the initial interaction between the SARS-CoV-2 N-protein and αS, multi-protein complexes are formed. In the presence of N-protein the onset of αS aggregation into amyloid fibrils is strongly accelerated, indicating that N-protein facilitates the formation of a critical nucleus for aggregation. Fibril formation is not only faster, but also proceeds in an unusual two-step process. In cells, the presence of N-protein changes the distribution of αS over different conformations that likely represent different functions at already short timescales. Disturbance of αS proteostasis might be a first step towards nucleation of fibrils. Our results point toward a direct interaction between N-protein of SARS-CoV-2 and αS as molecular basis for the observed relations between virus infections and parkinsonism. The observed molecular interactions thus suggest that SARS-CoV-2 infections may have long term implications and that caution is required in considering N-protein as an alternative target in vaccination strategies.

## Materials and Methods

### αS Production

Expression of recombinant human αS, the 140C mutant αS(140C) with a single alanine to cysteine substitution at residue 140 and the double cysteine mutant αS(9C/69C) was performed in E. coli B121 (DE3) using the pT7-7-based expression system. Details on the purification procedure are described elsewhere.^44^

### Preparation of labelled αS

To visualize the microinjected αS in cells, the αS(9C/69C) was labelled with a FRET pair as described before.^39^ In short the cysteines in the αS(9C/69C) were reduced with DDT. After removal of DDT, an equimolar concentration of the maleimide derivative of AF488 was added to 0.5 ml of 200 μM αS(9C/69C) and incubated for 1 hour at room temperature. To remove unreacted dye and DTT a Zeba Spin desalting column (Pierce Biotechnology) was used. The labelled protein was applied to a Thiopropyl Sepharose 6B column (GE Healthcare Life Sciences) to remove double labelled protein. Column bound single-labeled and/or unlabeled αS was eluted using 10-15 ml of 10 mM Tris-HCl, pH 7.4 buffer, containing β-mercaptoethanol. Subsequently a 2-3x molar excess of maleimide functionalized AF568 was added. After incubation for 1 hour at room temperature, free dye was removed using two desalting steps and the solution was filtered through a Microcon YM100 filter (Millipore, Bedford, MA). In the main text the FRET labelled αS(9C/69C) protein is referred to as the αS FRET probe. Single labelled αS was prepared using the αS140C cysteine mutant which was incubated with 2-3x molar excess of AF488 or AF647 maleimide. The free dye was removed using the protocol mentioned above.

### Aggregation assays

50 μM αS was aggregated in the presence of SARS-CoV-2 N-protein (PKSR030485, Elabscience, US) and S-protein (10549-CV-100, R&D systems, UK) at concentrations specified in the main text, 20 mM Tris buffer (Sigma-Aldrich, UK), pH = 7.4, 5 µM Thioflavin T (ThT, Fluka, Sigma-Aldrich, UK) and different concentrations of NaCl (Sigma-Aldrich, USA) as mentioned in the main text. The aggregations were performed in a 96-well half area clear flat bottom polystyrene NBS (low bind) microplate (3881, Corning, US) while shaking at 500 RPM and 37 °C. To follow the formation of amyloid fibrils the increase of ThT fluorescence was monitored using a plate reader (Infinite 200 Pro, Tecan Ltd., Switzerland). The ThT dye was excited at 446 nm and the fluorescence signal was measured at 485 nm every 10 minutes. Samples were prepared in 3-5 replicates of 50 μl. The lag time is defined as the time point at which a twofold increase in the fluorescence intensity compared to the initial intensity values (baseline) is observed.

### Microscale thermophoresis

The binding between αS and N-protein was studied using a Monolith NT.115 (NanoTemperTechnologies GmbH, Germany) MST system. The thermophoretic movement was monitored at a constant 20 nM concentration of αS140C-AF488, and a dilution series of concentrations of N-protein. Samples were prepared in 20 mM Tris-HCl, pH 7.4, 10 mM NaCl and transferred to capillaries (Standard treated, NanoTemper Technologies GmbH, Germany), measurements were performed at 37 °C with a constant blue LED power of 20% and at MST infrared laser powers of 40 % to induce thermophoretic motion. For each capillary the infrared laser was switched on 10 s after the start of the measurement for a 30 s period, followed by another 10 s period with a turned off infrared laser to record the back diffusion. Data were analyzed with the MO.Affinity Analysis software.

### Fluorescence microscopy and fluorescence correlation spectroscopy

Fluorescence images were obtained on a laser scanning confocal microscope (MicroTime 200 with a FlimBee scanner, PicioQuant, Germany). To excite either DAPI or FRET labelled αS sequential 405nm and 485nm laser excitation in combination with a dichroic mirror (ZT405/485rpc-UF3, Chroma, USA) was used. A UPlanSApo, 60x, 1.2 NA objective (Olympus, Japan) was used for imaging. Emission was detected with Single Photon Counting Modules (SPCM-AQRH-15, Excelitas, Canada). DAPI emission was detected via a bandpass filter (F02-447/60-25, Semrock, USA), emission from the FRET pair via a 488 long pass emission filter (LP02-488RU-25, Semrock, USA). The FRET signal was further split by a 585 nm dichroic beam splitter (T585lpxr, Chroma, USA) into a green channel (bandpass FF01-520/35-25, Semrock, USA) for detection of the FRET-donor signal and a red channel (bandpass BA590, Olympus, Japan) for detection of the FRET-acceptor signal. Fluorescence intensity images were exported as raw data and used for image analysis and visualization purposes.

The fluorescence correlation spectroscopy (FCS) experiments were performed on the same setup. For the FCS experiments samples were prepared containing 10 nM αS140C-AF647, 10 nM αS (total of 20 nM αS) and a range of N-protein concentrations in 20 mM Tris buffe, pH 7.4. To excite AF647 labelled αS, a 640nm laser was used. Excitation light was reflected towards the sample and emission was separated from excitation light using a dichroic mirror (ZT488/640rpc-UF3, Chroma, USA). The emission was further filtered spectrally by a 690 nm bandpass. Before detection the emission was filtered spatially by a 100 µm pinhole. Autocorrelation curves were calculated and analyzed using the SymPhoTime 64 software. A diffusion model that includes the triplet state population was used to fit the data. For each experimental condition, 10 measurements with duration of 10 seconds were recorded and analyzed.

### Atomic force microscopy

A 10 μl volume of 5 times diluted aggregated sample (initial αS concentration = 50 µM and N-protein concentration = 0.2 – 1 μM) was deposited onto freshly cleaved mica (Muscovite mica, V-1quality, EMS, US) and left to rest for 5 min. Then the sample was carefully washed 4 times with 20 μl of demineralized water (MilliQ) and gently dried under a low flow of nitrogen gas. AFM images were acquired using a BioScope Catalyst (Bruker, US) in soft tapping mode using a silicon probe, NSC36 tip B with a force constant of 1.75 N/m (MikroMasch, Bulgaria). Images were captured with a resolution of 512 × 512 (10 μm x 10 μm) pixels per image at a scan rate of 0.2 to 0.5 Hz. AFM images were processed with the Scanning Probe Image Processor (SPIP, Image Metrology, Denmark) and the Nanoscope Analysis (Bruker, Us) packages. Fibril morphology was analyzed using a custom fibril analysis Matlab script adapted from the FiberApp package.^45^

### Fluorescence spectroscopy

A Varian Eclipse spectrofluorometer was used to record bulk emission spectra of the FRET labelled αS(9C/69C) (500nM) and SARS CoV-2 N-protein (5 µM) mixtures in 1xPBS buffer. The samples were pipetted into a 3 mm quartz cuvette and excited at 488nm wavelength, and emission was collected from 500 nm to 700 nm, with a slit width of 5 nm both for excitation and emission.

### Cell culture, microinjection, and nuclear counterstaining

SH-SY5Y cells (ATCC, USA) were grown in proliferation medium (DMEM-F12 GlutaMAX™ + 10% heat inactivated FBS + 1% non-essential amino acids +10 mM HEPES buffer + 1% Penicillin/Streptomycin; Gibco^®^, Invitrogen, USA). For microinjection the cells were seeded on glass bottom µ-Dishes with a cell location grid (grid-50, 35 mm, ibidi^®^, Germany). Dishes in which the cell confluency reached 70% to 90% were used for microinjection.

The microinjections were performed as described before.^39^ In short, the microinjections were performed using a FemtoJet^®^ (Eppendorf, Germany), equipped with a manual hydraulic 3D micromanipulator (Narishige, Japan). For injection, UV-O_3_ cleaned glass micropipettes with an inner diameter of approximately 400 nm (WPI, USA) were used. For microinjection the following concentrations of proteins in PBS were used, 500 nM FRET labeled αS(9C/69C) for 5 days incubation, 500 nM FRET labeled αS(9C/69C) and 20 µM αS for 3 days incubation, with and without 500 nM N-protein. The prepared solutions were used to backfill the glass micropipettes. For injection the following settings were used: an injection pressure of 150 hPa, a constant pressure of 15 hPa, and a duration of the injection of 0.1 second. During injections the sample was observed on a Nikon TE2000 microscope (Nikon, Japan).

After microinjection, the cells were incubated for either 3 or 5 days. Subsequently the samples were washed with PBS (3x) and fixed in 3.7% paraformaldehyde/PBS solution for 10 minutes at room temperature followed by an additional washing step with PBS. The cell nuclei were counterstained with 4′,6-diamidino-2-phenylindole (DAPI) at a final concentration of 300 nM for 10 minutes.

### Analysis of cell images

To estimate the FRET efficiency we first applied a threshold to the recorded fluorescence intensity images. Only intensities exceeding the mean intensity + 3x SD of each image were taken into account for the analysis of the FRET efficiencies. This thresholding was used to exclude background and noise from the analysis. From the thresholded FRET donor and acceptor fluorescence images, the E_FRET_ value per pixel was estimated as the ratio between intensity in the FRET acceptor channel divided by the sum of the intensities in the FRET donor and FRET acceptor channels. Note that these FRET efficiency values are estimates. We did not correct for acceptor cross excitation and donor bleed through in the acceptor channel since we only focus on differences within an image and changes between conditions. The mean FRET efficiency of all pixels in an image is represented as a single count in the cumulative histograms.

## Acknowledgements

We would like to thank Kirsten A. van Leijenhorst-Groener for the production of the recombinant α-synuclein protein. We are grateful to the Dutch Parkinson’s disease foundation “Stichting Parkinson-Fonds” for financial support.

## Author Contributions

C.B., I.S.N. and M.M.A.E.C. designed research; M.A.A.F and S.A.S performed the research; S.A.S. and M.A.A.F. analyzed and interpreted the data; all authors contributed to writing the paper.

