## Supporting information for "Interactions between SARS-CoV-2 N-protein and α-synuclein accelerate amyloid formation"

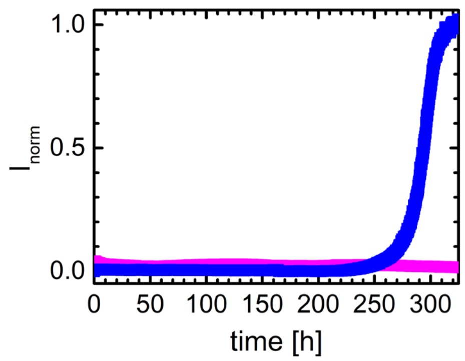


**Figure S1**. ThioflavinT (ThT) assay. Even at the highest concentrations used in our studies the incubation of N-protein alone (1 µM, magenta) does not result in an increase in the ThT fluorescence. For comparison we show the increase in ThT fluorescence upon aggregation of 50 µM αS (blue) as used in our aggregation studies.

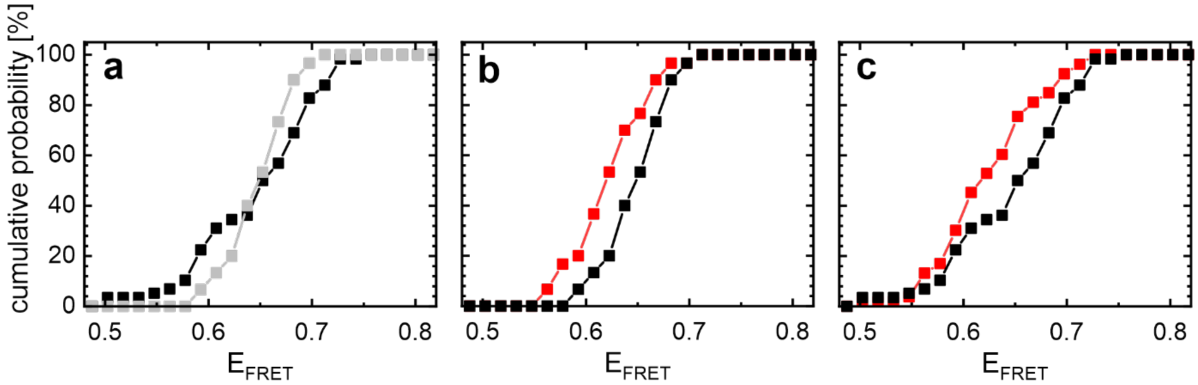


**Figure S2**. Cumulative histograms for the averaged αS FRET efficiency per image ($E_{FRET}$). **a)** For the control samples, containing no N-protein, the $E_{FRET}$ distributions obtained 5 days (black) and 3 days and additional αS (gray) after injection agree well. The width of the distribution is slightly enhanced for the control data obtained 5 days after microinjection. The mean $E_{FRET}$ value quantitatively agrees between the two sets of control experiments. b) Cumulative histograms for the averaged αS FRET efficiency per image for the N-protein injected cells (red) and the control cells (black) 3 days after injection. The distribution of the αS FRET efficiencies for the N-protein injected cells is shifted to lower values compared to the control. c) As b) but now for cells 5 days after microinjection, without additional αS.

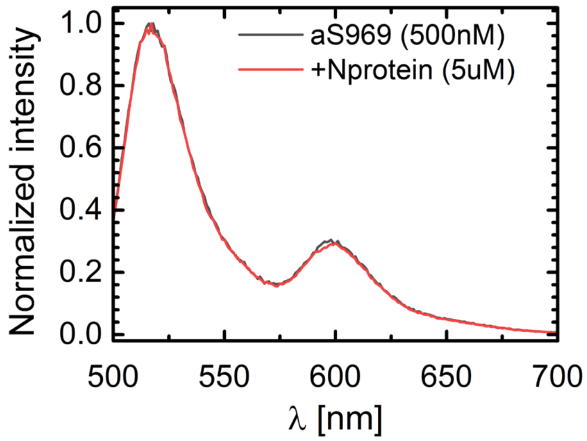


**Figure S3**. Fluorescence spectra of the αS(9C/69C) FRET probe. The spectrum in black shows the emission of the FRET probe alone at a concentration of 500 nM. The red spectrum shows the emission spectrum of the FRET probe (500nM) in the presence of excess N-protein (5 µM). Although N-protein and αS interact as shown in Figure 2 of the main text, this interaction does not lead to a change in the energy transfer process between the donor and acceptor fluorophore. From this we conclude that the N-protein does not interact with αS in the FRET labelled region, alternatively binding does not induce measurable conformational changes in αS.
